# Diverse tumorigenic consequences of human papillomavirus integration in primary oropharyngeal cancers

**DOI:** 10.1101/2021.06.20.449168

**Authors:** David E. Symer, Keiko Akagi, Heather M. Geiger, Yang Song, Gaiyun Li, Anne-Katrin Emde, Weihong Xiao, Bo Jiang, Jingfeng Li, Amit D. Agrawal, Enver Ozer, Adel El Naggar, Zoe Du, Jitesh Shewale, Birgit Stache-Crain, Mark Zucker, Nicolas Robine, Kevin R. Coombes, Maura L. Gillison

## Abstract

Human papillomavirus (HPV) causes 5% of all cancers and frequently integrates into host chromosomes, but the impacts of integration in tumorigenesis remain unclear. Analysis of 105 HPV-positive oropharyngeal cancers by whole genome sequencing detects viral integration in 77%, revealing five statistically significant integration hotspots near genes that regulate epithelial stem cell maintenance (i.e. *SOX2, TP63, FGFR, MYC*) and immune evasion (i.e. *CD274*). Somatic hyperamplification is enriched 16-fold near HPV integrants, and the extent of focal host genomic instability increases with local density of HPV integrants. Genes expressed at extreme outlier levels are increased 86-fold within +/- 150 kb of integrants. Across 95% of tumors with integration, host gene transcription is disrupted via intragenic integrants, chimeric transcription, outlier expression, gene breaking and/or de novo expression of noncoding or imprinted genes. We conclude that HPV integration contributes substantively to cancer development by causing extensive disruption of host genome structure and gene expression.

## INTRODUCTION

Human papillomavirus (HPV) infection causes approximately 5% of all human cancers, resulting in 650,000 cases worldwide each year (de Martel et al. 2017). These include anogenital cancers and a subset of oropharyngeal cancers that is increasing markedly in incidence (Tota et al. 2019). The HPV oncoproteins E6 and E7 promote host genomic instability in multiple ways including degradation of tumor protein p53 (TP53) and RB transcriptional corepressor 1 (RB1), respectively. E6 and E7 expression is necessary but not sufficient for HPV-associated carcinogenesis. Secondary genetic events such as host gene mutations also are required (Cancer Genome Atlas Research et al. 2017; Gillison et al. 2019).

HPV integration into cervical cancer genomes was first reported over 30 years ago (Durst et al. 1987). Subsequent studies were unable to reveal the full extent of HPV integration because they utilized biased and/or insensitive laboratory techniques to detect and map integrants (e.g. Southern blotting, PCR, whole exome sequencing [WES], RNA sequencing [RNA-seq] and others) (Cancer Genome Atlas Research et al. 2017), reviewed in (Bodelon et al. 2016). More recently, a hybrid capture-based method was used to report recurrent hotspots of HPV integration in cervical dysplasias and cancers (Hu et al. 2015). However, virus-host breakpoints were identical at a single nucleotide level across multiple samples, raising questions about artifacts and undermining the data and conclusions (Dyer et al. 2016). Recent TCGA genomic studies of oropharyngeal and cervical cancers mostly used RNA-seq to map integration sites indirectly, and subsets studied with whole genome sequencing (WGS) were too small to identify recurrent hotspots (Parfenov et al. 2014) (Cancer Genome Atlas Research et al. 2017). In sum, the impacts of HPV integration on host genome structures and gene expression have yet to be defined and quantified comprehensively across an adequately powered collection of tumors.

Using WGS to study relatively small numbers of HPV-positive cancers and cell lines, we and others have shown that HPV integrants recurrently flank or bridge focal regions of extensive host genomic instability, including copy number variation (CNV) and structural variation (SV) (Akagi et al. 2014)(Parfenov et al. 2014) (Cancer Genome Atlas Research et al. 2017). We proposed a mechanistic, looping model by which replication of transient, circular, virus-host intermediate structures (using the HPV origin of replication) is followed by recombination and repair, leading to integrated HPV-host concatemers and extensive genomic structural variation (Akagi et al. 2014). We hypothesize that host genomic alterations associated with HPV integration are critical contributors in the pathogenesis of a majority of HPV-positive primary cancers. Here we report a comprehensive analysis of virus integration and its impacts in 105 HPV-positive oropharyngeal squamous cell carcinomas (OPSCC) that is powered to detect statistically significant recurrent hotspots of HPV integration, using precise and unbiased genomics methods including WGS and RNA sequencing (RNA-seq).

## RESULTS

### Genomic sites of HPV integration reveal clustering in individual tumors

WGS data from tumor and normal blood leukocyte (T/N) pairs from 105 patients with newly diagnosed HPV-positive OPSCC reveal that a majority (88%, n=92) harbor HPV16 (Gillison et al. 2019). Overall, 874 virus-host breakpoints were identified across 81 (77%) tumors, based on detection of split and/or discordant sequencing reads mapping both to HPV and the reference human genome (**Fig. S1.1**). As we observed in cultured HPV-positive cancer cells (Akagi et al. 2014), HPV breakpoints in primary tumors are uniformly distributed across the entire ~8 kb viral genome, and thus are not enriched preferentially in any particular viral gene (**Fig. 1A, Fig. S1.2**). These data refute an accepted paradigm about preferential disruption of HPV E2 by insertional breakpoints, which in turn would dysregulate HPV E6 and E7 oncogene expression and promote carcinogenesis (Romanczuk and Howley 1992).

**Fig. 1.**
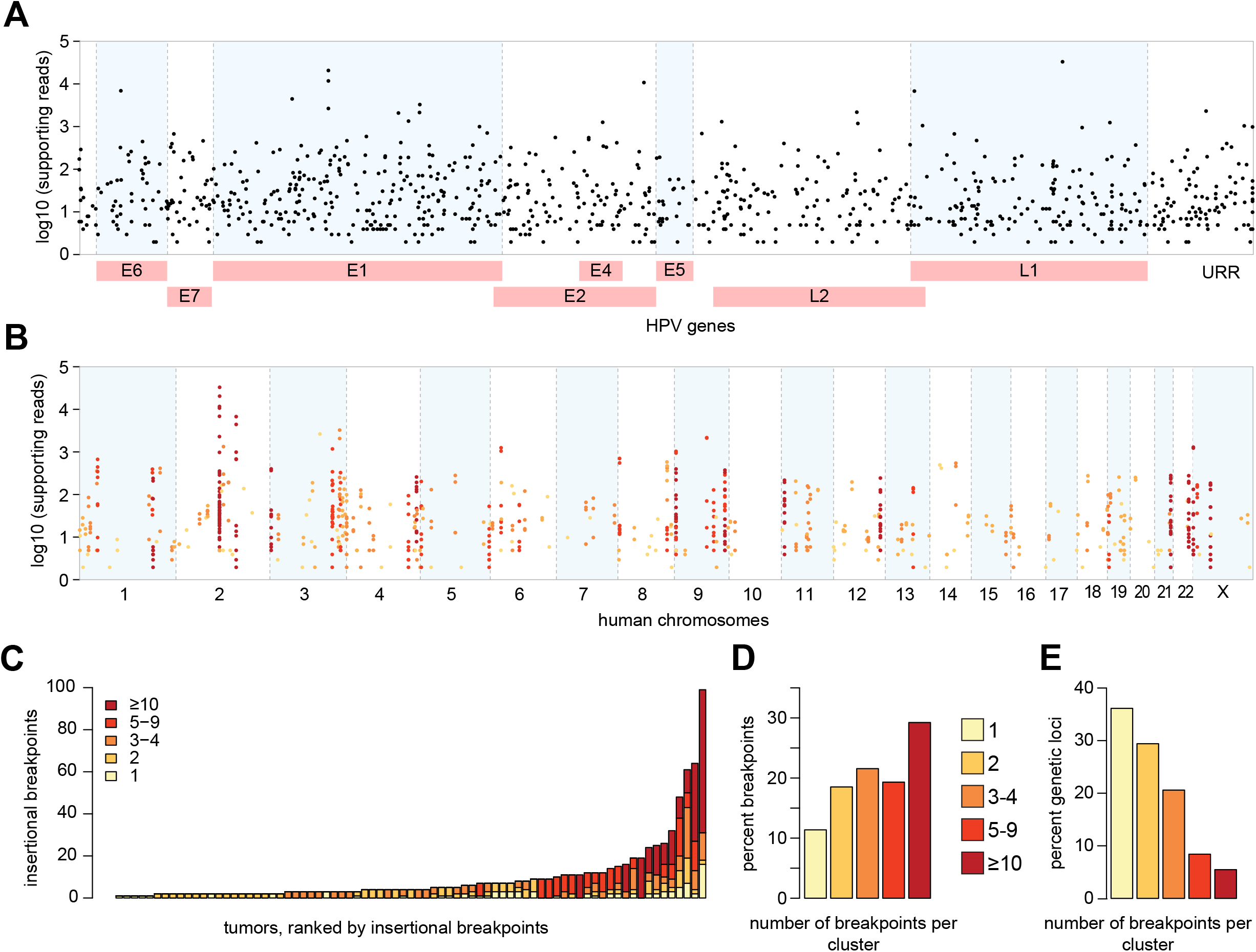
Frequent clustering of virus-host breakpoints in individual tumors. (A) Breakpoints (*dots*, n = 874) identified in 105 HPV-positive OPSCCs mapped to HPV16 genome (*x-axis*). Non-HPV16 breakpoint (n=50) coordinates are approximated. *Y-axis*, log10 of WGS reads supporting each breakpoint. (B) Breakpoints uniquely mapped to the human genome (*x-axis*, n = 756, hg19) are clustered within 500 kb windows. *Y-axis*, as per (A). *Colors*, breakpoints in individual tumors in clusters, as per key in panel C. (C) Counts of uniquely mapped breakpoints (*y-axis*), ranked by frequency in individual HPV-positive OPSCCs (*x-axis*). *Key, colors*, breakpoint counts per cluster. (D) Overall frequencies of breakpoints in clusters across all tumors (*x-axis, colors*, breakpoint counts per cluster). (E) Overall frequencies of distinct genetic loci harboring clusters of various breakpoint counts within 500 kb (*x-axis, colors*) in individual tumors. See also **Table S1**, **Fig. S1**.

Of the 874 breakpoints, 756 (86.5%) map uniquely in 77 (73%) tumors, while the remainder align either to repetitive elements (n=95) or unassigned contigs (n=23, **Table S1.1**). No breakpoints are detected in 24 (23%) tumors, indicating that they harbor episomal virus only. This confirms that virus integration is not mandatory for HPV-associated cancer development. Across the tumors, uniquely mapped breakpoints are broadly distributed across the human genome **(Fig. 1B, Table S1.2**). Although median viral copy numbers are similar in tumors with integrated vs. exclusively episomal HPV, the variance in copy numbers is greater across the former (**Fig. S1.3**). In contrast with prior reports (Koneva et al. 2018), we find no significant association between integration status and patients’ demographic characteristics or clinical outcomes (**Fig. S1.4, Table S1.3**).

We used several independent methods to validate subsets of uniquely mapped breakpoints. HPV capture-seq, utilizing custom HPV16 baits, confirms 92% of breakpoints detected by WGS in three HPV-positive OPSCC (**Table S1.4**, and below). We previously used Sanger sequencing to confirm 95% of breakpoints detected in cell lines (Akagi et al. 2014), so we randomly selected ~10% of breakpoints here for similar validation. Sanger sequencing confirms 100% of custom PCR amplicons, revealing microhomology between virus and host sequences occurring significantly more than expected by chance at breakpoints (**Fig. S1.5, Table S1.5**) (Symer et al. 2002). We also detect insertions of heterologous or untemplated DNA sequences, but find no identical breakpoint sequences across samples (Hu et al. 2015; Dyer et al. 2016). These data support contributory but nonessential roles for nonhomologous end-joining (NHEJ) and/or microhomology-mediated end joining (MMEJ) as DNA repair mechanisms in HPV integration (Hu et al. 2015; Leeman et al. 2019).

Mapping of all detected breakpoints against the human genome prompted an initial interpretation that integrants occur at genomic hotspots (**Fig. 1B**). However, closer inspection reveals breakpoint clustering within individual tumors. We define a cluster as three or more unique breakpoints within a 500 kb genomic segment in a single tumor (**Fig. 1B and 1C**). We and others have reported comparable breakpoint clusters in HPV-positive cell lines derived from cervical (e.g. HeLa) and oropharyngeal (e.g. UPCI:SCC090 and others) cancers (Adey et al. 2013; Akagi et al. 2014). Breakpoint counts per tumor vary widely (mean 9, median 4, range 1-99) (**Fig. 1C**). Of 756 uniquely mapped breakpoints, 70% are located within a cluster in an individual tumor (**Fig. 1D**), with a median of one cluster per tumor (range 0 to 9). By contrast, when considering all 500 kb loci harboring at least one breakpoint, the majority contains simple insertions rather than clusters (**Fig. 1E**).

### Integration hotspots target genes involved in epithelial stem cell maintenance and immune evasion

Upon accounting for clustered integrants in individual tumors, we identified several distinct genetic loci at which breakpoints are detected across three or more independent tumors, implicating recurrent genomic hotspots for HPV integration. Null hypothesis testing, involving a model of targeting megabasepair (Mbp) genomic segments across the 105 WGS tumors, identifies five statistically significant, distinct hotspots of recurrent integration, including *SOX2, TP63, FGFR3, MYC* and *CD274* (**Fig. 2A, Table S2.1**). Each of these genes has well-established roles in epithelial stem cell maintenance or anti-tumor immunity. A separate, gene-centric bioinformatics approach confirms that the same 5 genes are recurrently targeted hotspots (**Table S2.2**). Both the *MYC* and *TP63* loci also were identified as genomic hotspots in cervical cancers (Bodelon et al. 2016).

**Fig. 2.**
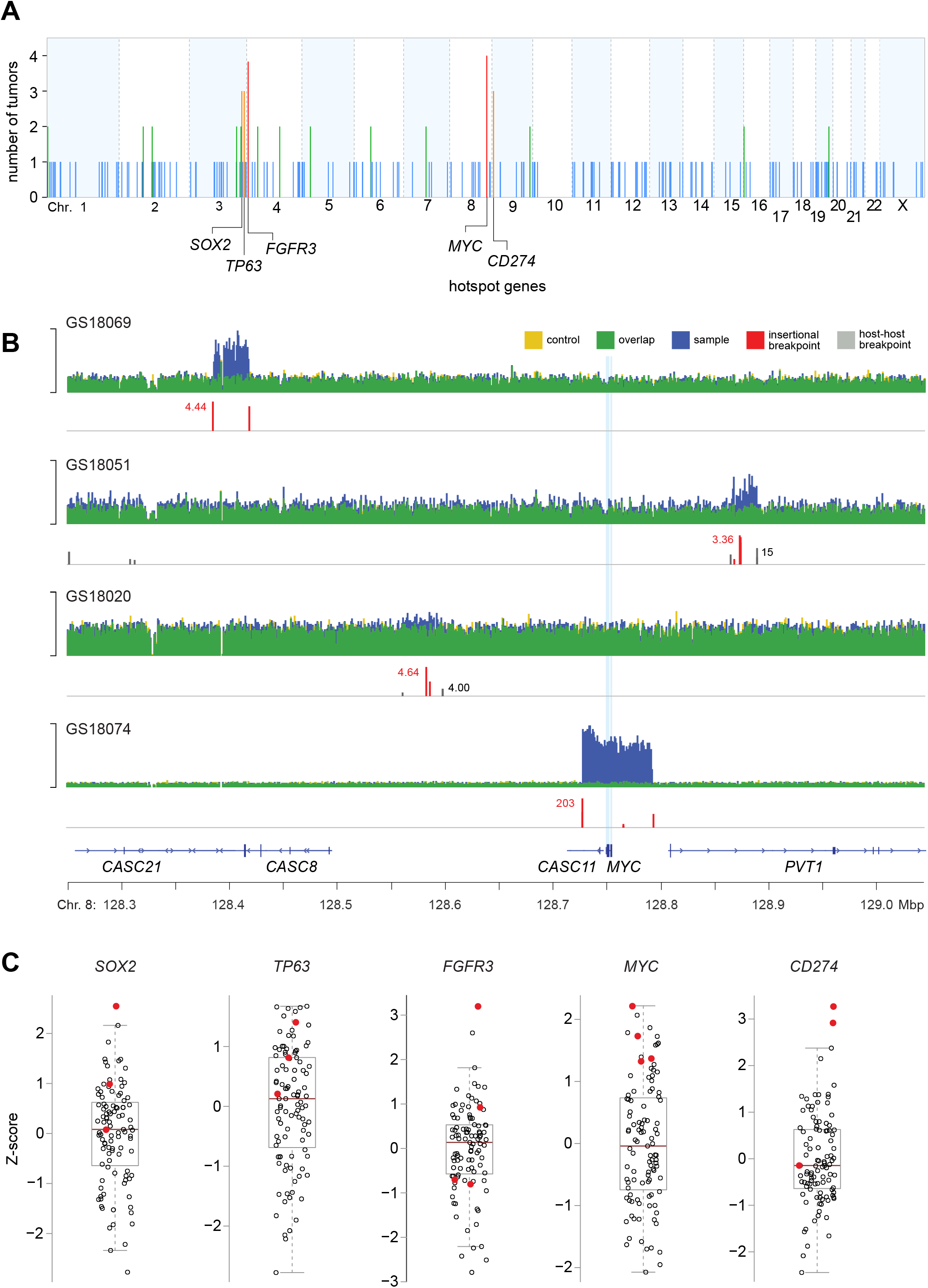
Genomic hotspots of integration near genes involved in epithelial stem cell maintenance and immune evasion. (A) Counts of independent tumors harboring ≥1 virus-host breakpoints (*y-axis*) in 1 Mbp genomic segments across the human genome (*x-axis*). Recurrent hotspots (*orange*, n=3 tumors; *red*, n=4) at segments containing *SOX2, TP63, FGFR3, MYC*, and *CD274* (empirical probability, p=1E-6). Other integration sites are not statistically significant hotspots in these tumors (*blue*, n=1 tumor; *green*, n=2). (B) IGV browser views of WGS depth of coverage (*y-axis*) and virus-host breakpoints (*red*) in 4 independent tumors within a 1 Mbp genomic segment containing *MYC* (*x-axis*; *light blue vertical lines*, exons). *Colors* as in *key, top*. (C) Transcript levels of *SOX2, TP63, FGFR3, MYC*, and *CD274 (y-axis*, Z-score of log2 TPM value), in tumors quantified by RNA-seq (*circles*, n=103). *Red fill*, tumors with breakpoints near the hotspot genes (panels A, B). *Box and whiskers*, median (*brown horizontal line*), quartiles (*light gray box*). See also **Table S2**, **Fig. S2**.

Within the one Mbp genomic segment containing *MYC*, breakpoints were identified in four tumors and are associated with flanking CNVs and SVs (**Fig. 2B**). In one case, breakpoints directly flank a ~9-fold amplification of *MYC*, and in another they flank a 3-fold amplification upstream of *MYC*. Both are associated with elevated *MYC* transcript levels (**Fig. 2C**), consistent with impacts of amplification of the gene or adjacent super-enhancers (Zhang et al. 2016). HPV integration near *MYC* previously was reported in cervical cancers and derived cell lines (Peter et al. 2006; Yuan et al. 2017).

Additional examples of integration hotspots targeting “stemness” genes involve *FGFR3, SOX2* and *TP63*. In four tumors, breakpoints map to a hotspot containing *FGFR3* (**Fig. 2C, Fig. S2.1**). We previously documented recurrent FGFR3 p.249S>C activating mutations in 11% of HPV-positive OPSCCs (Gillison et al. 2019). Cancer stem cell renewal is sustained by FGF pathway activation with downstream regulation of several key transcription factors (Mossahebi-Mohammadi et al. 2020). Another hotspot involves *SOX2*, encoding a stem cell pluripotency factor. In one tumor, insertional breakpoints flank ~5-fold amplification with outlier transcript upregulation (**Fig. 2C, Fig. S2.1**). Such SOX2 overexpression promotes proliferation and anchorage-independent growth of squamous epithelial cancers (Bass et al. 2009). Hotspot integrants also map near *TP63*, encoding a transcription factor that regulates squamous epithelial stem cell maintenance, differentiation and proliferation (Senoo et al. 2007). Our prior analysis identified inactivating mutations in genes promoting epithelial differentiation (i.e. *ZNF750, KMT2D, RIPK4* and *TGF-beta*) in 37% of HPV-positive OPSCC (Gillison et al. 2019). We conclude that disruption of differentiation and maintenance of epithelial stemness are important in the pathogenesis of these cancers.

A recurrent integration hotspot in three tumors involves the immune checkpoint ligand gene *CD274*, encoding programmed cell death 1 ligand 1 (PD-L1). HPV integration near this gene has been reported previously (Cancer Genome Atlas Research et al. 2017; Koneva et al. 2018). In two of the cases studied here, HPV integrants are associated with 5-10 fold amplification and outlier expression of *CD274* (**Fig. 2C, Fig. S2.1**). In a tumor with 63 breakpoints, clusters are identified on Chr. 4, 5, 9, 10, 19 and 22 in direct proximity to CNVs affecting specific host genes (e.g. *CD274* and *EP300*, **Fig. 3A**, **Fig. S3.1**). Inspection of WGS depth of coverage around a cluster of 18 breakpoints on Chr. 9p.24.1 reveals extensive CNV and SV, including ~11-fold amplification of *CD274* **(Fig. 3A)**. Linked-read sequencing (10X Genomics) demonstrates that the HPV integrants co-occur with genomic deletions and amplifications on the same haplotype (**Fig. 3B, Fig. S3.2-S3.3**). Barcodes of linked reads mapping to the CNVs or SVs are shared at high frequencies with HPV16, establishing direct connectivity between integrants and flanking structural variants. In these cases, HPV integration near *CD274* likely promoted immune escape and tumor development.

**Fig. 3.**
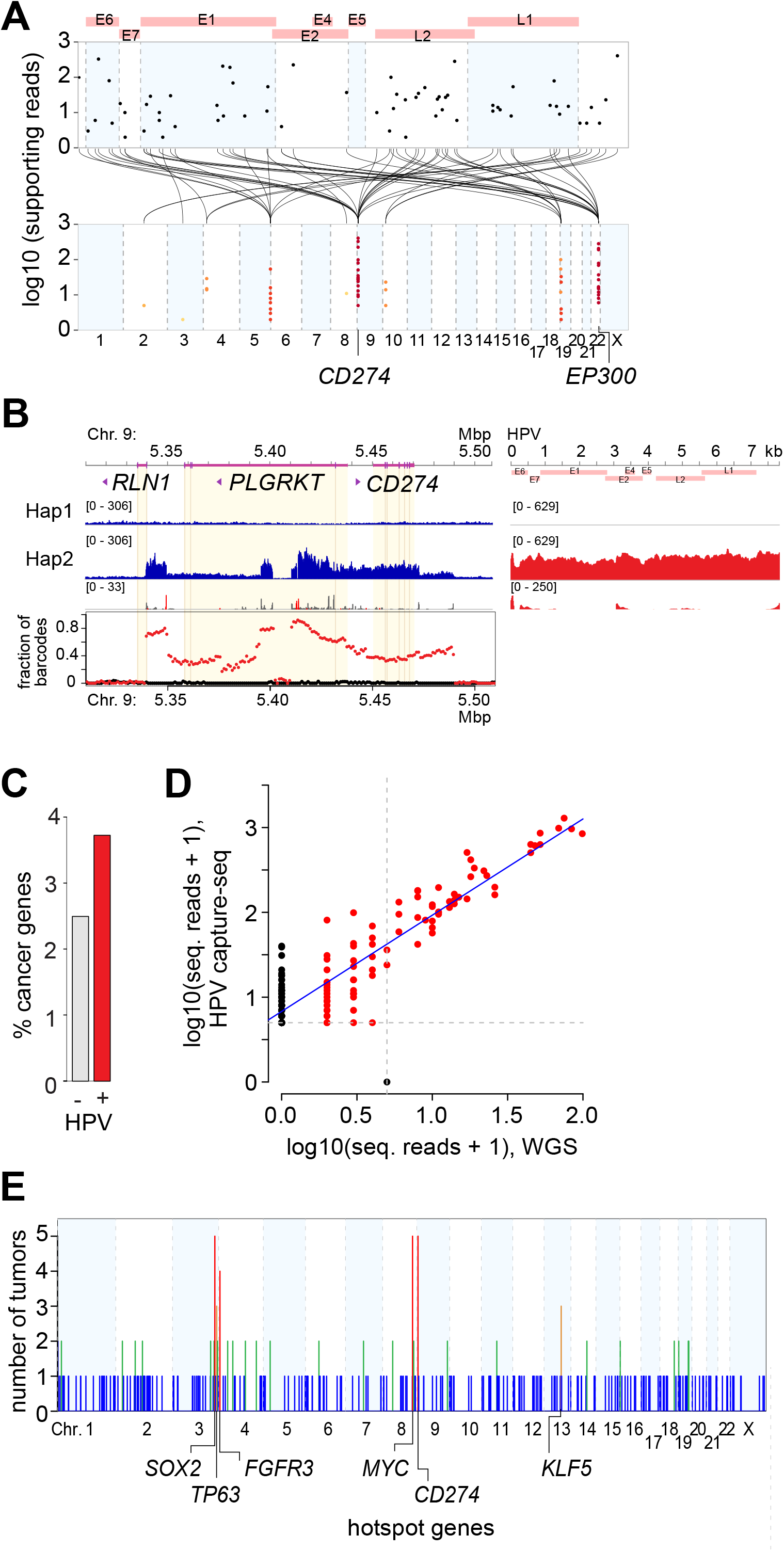
Associations between HPV integrants, CNVs and SVs in individual tumors. (A) Strudel plot shows virus-host breakpoints in a representative OPSCC, GS18047. Breakpoints mapped to the HPV16 genome (*top, x-axis*) are connected (*black lines*) with the host genome (*bottom, x-axis*), clustered on Chrs. 4, 5, 9, 10, 19 and 22 (*colored dots, as per key, Fig. 1C*). Disrupted genes include *CD274* and *EP300*. (B) Haplotype-resolved linked reads (*blue*, host depths of sequencing coverage; *red*, HPV) connect HPV16 sequences (*right*), virus-host breakpoints (*red peaks*) and host-host breakpoints (*gray*) on one allele (haplotype 2) at the *CD274* locus on Chr. 9p24.1. *Graph, bottom*, shared linked-read barcodes connect HPV16 exclusively to haplotype 2 (*red*) but not haplotype 1 (*black*). (C) Fraction of genes with (*red*) or without (*gray*) HPV breakpoints within +/- 500 kb that are annotated cancer genes (*y-axis*) (Sondka et al. 2018). Fisher’s exact, p=6.3E-5. (D) Scatterplot shows strong correlation between read counts supporting individual breakpoints (*red dots*) identified with HPV capture-seq (*y-axis*, n=164) vs. WGS (*x-axis*, n=86) in the same tumor (r=0.91; p=1.8E-63). (E) Adding 53 tumors studied by HPV capture-seq to 105 tumors studied by WGS (**Fig. 1A**), tumors harboring ≥1 virus-host breakpoints (*y-axis*) in 1 Mbp genomic segments were recounted across the human genome (*x-axis*). Statistically significant, recurrent hotspots (*orange*, n=3 tumors; *red*, n=4 or 5) are detected at segments containing *SOX2, TP63, FGFR3, MYC, CD274* and *KLF5* (empirical probability, p=7E-6). See also **Fig. 2, Table S3, Fig. S3**.

Cancer-driving genes (Sondka et al. 2018) are enriched in genetic loci neighboring HPV insertional breakpoints over loci that lack such breakpoints (**Fig. 3C**). Ontology analysis of genes in these loci reveals enrichment of genes involved in regulation of activated T cell proliferation, somatic stem cell maintenance, and mitochondrial apoptotic signaling, among others (**Table S3.1**). These findings further confirm enrichment of HPV integrants near genes and pathways involved in cancers, and strongly implicate viral integration as a driver of carcinogenesis by clonal selection.

### HPV capture-seq data from additional tumors identify another hotspot involving epithelial stemness genes

To confirm and extend the recurrent HPV integration hotspots identified from WGS data (**Fig. 2**), we analyzed additional HPV-positive OPSCC with HPV capture-seq, a targeted sequencing method (Warburton et al. 2018). First, we compared insertional breakpoints identified from HPV capture-seq vs. WGS from the same tumor. Upon normalization of sequencing depth of coverage, numbers of supporting reads are closely correlated for breakpoints detected by both methods (**Fig. 3D**).

Based on these results, we used HPV capture-seq to identify HPV integrants in 53 additional HPV-positive OPSCC (**Table S3.2 – S3.3**). These independent tumors harbor breakpoints near the same hotspot genes noted above, adding further support for them (i.e. at *SOX2, MYC, CD274*). By combining these 53 tumors with the 105 WGS tumors, we identify an additional, significant hotspot in three tumors near the zinc-finger transcription factor Kruppel-like factor 5 (*KLF5*) on chr13q22.1 (**Fig. 3E**, **Table S3.4 – S3.5**). *KLF5* is a candidate oncogene that regulates stemness, proliferation and differentiation of the basal epithelial cell (Ghaleb et al. 2005), the cell specifically infected by HPV. We conclude that HPV integration near genes that regulate epithelial stem cell maintenance confers a selective growth advantage, promoting tumorigenesis.

### Enrichment of HPV integrants in genomic regions with CNVs and SVs

Breakpoint clusters in individual tumors with identified hotspots frequently are associated with CNVs and SVs (**Figs. 2 and 3**). Therefore, we investigated amssociations between HPV integration and CNVs and SVs across all tumors studied by WGS. The frequency distribution of CNVs is markedly different in comparing 100 kb host genomic segments with and without virus-host breakpoints (**Fig. 4A**). Quantile-quantile (Q-Q) plots comparing the distribution of genomic copy numbers in the presence vs. absence of HPV breakpoints demonstrate unequivocally that viral insertions are strongly associated with copy number alterations (**Fig. 4B**). Breakpoints are highly enriched in segments containing CNVs, particularly in hyper-amplified segments with estimated ploidy ≥ 4N (16.3-fold enrichment, binomial test, adj. p=2.06E-20, **Table S4)**. Breakpoints directly flank CNV regions with amplification up to 15-fold and/or lengths exceeding 5 Mbp.

**Fig. 4.**
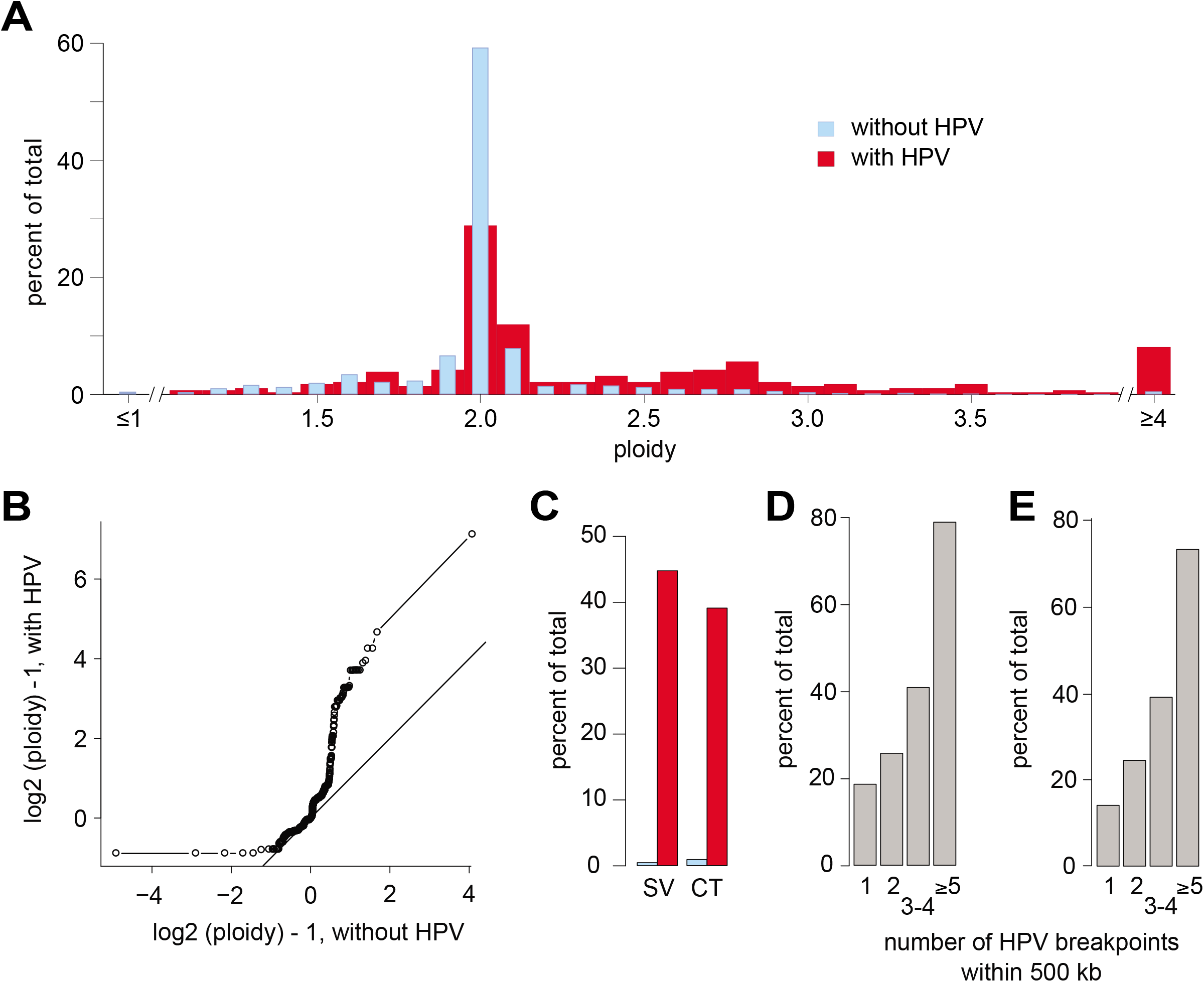
HPV integrants are associated with CNVs and SVs across tumors. (A) Shown are distinct frequency distributions (*y-axis*) of ploidies (*x-axis*) of 100 kb genomic segments with (*red*) vs. without (*blue*) virus-host breakpoints across 105 HPV-positive OPSCC (chi-squared, p=1.8E-18). (B) Quantile-quantile (Q-Q) plot confirms differences in copy numbers of genomic segments with (*y-axis*) and without (*x-axis*) breakpoints, deviating significantly from the line of identity. (C) Frequencies (*y-axis*) of structural variation (SV, *left*) and step-changes in copy number (copy number transition [CT] +/-0.5N, *right*) are significantly greater in 100 kb segments with a breakpoint (*red*) vs. without (*gray*) (SV, binomial test, one-tailed, p=3.3E-224; CT, p=2.39E-14). (D, E) Among 500 kb genomic segments with ≥1 breakpoints, frequencies (*y-axis*) of (D) SVs and (E) CTs increase with breakpoint counts in the cluster. See also **Table S4**.

Genomic SVs including deletions, insertions, inversions and chromosomal translocations are enriched in genomic segments with HPV breakpoints compared to those without (44.7% vs. 0.47%, **Fig. 4C**). Copy number transitions (CT, i.e. step changes in ploidy >0.5 N) also are enriched in segments with breakpoints compared to those without (37.6% vs. 0.88%; **Fig. 4C**). Across all tumors, breakpoints map within 10 kb of inversions in 21%, duplications in 34%, deletions in 20%, and chromosomal translocations in 6%. The larger the number of breakpoints within a cluster, the more frequent the concomitant SV counts and step-changes in CNVs (**Fig. 4D and E**), supporting direct involvement of HPV integration in generation of host genomic rearrangements.

### Association between HPV integrants and outlier expression of neighboring host genes including cancer genes

Z-scores were calculated for all genes’ transcript expression levels across 103 OPSCC tumors with available RNA-Seq data. To investigate impacts of HPV integration on host gene expression, we identified neighboring genes within +/-500 kb of breakpoints in affected tumors, and used Q-Q plots to compare the distribution of expression levels for these genes near a breakpoint vs. those without a corresponding breakpoint. In many cases with breakpoint(s) present, neighboring genes are significantly overexpressed (and less frequently underexpressed) in the affected tumor compared to controls, as shown by significant deviation of many data points away from the line of identity (**Fig. 5A**).

**Fig. 5.**
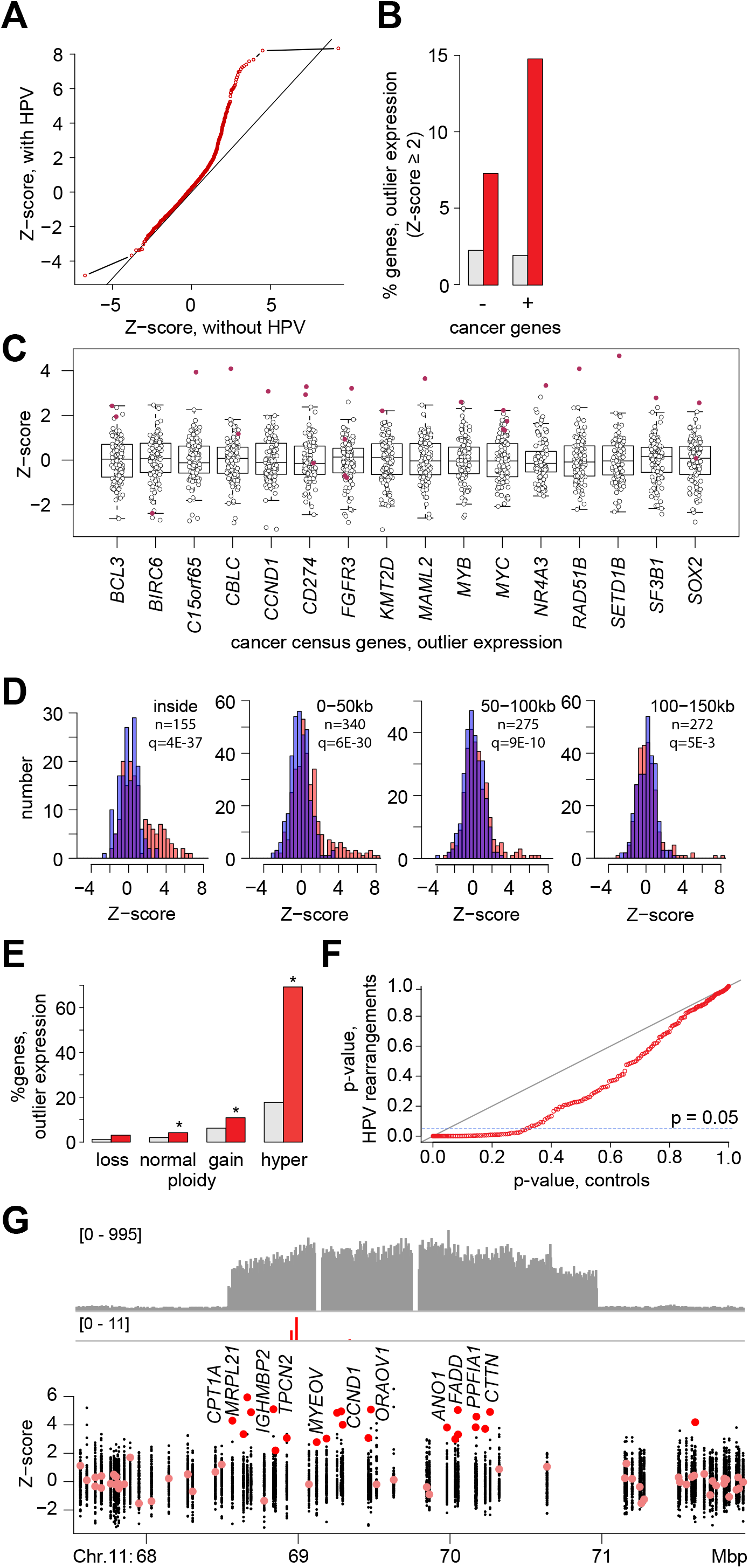
HPV integrants are associated with outlier expression of neighboring host genes. (A) Q-Q plot compares Z-score distributions of expression levels for genes near (+/- 500 kb) virus-host breakpoints (*y-axis*) vs. expression of the same genes without nearby breakpoints in all other tumors (*x-axis*; Kolmogorov-Smirnov test, p< 2.2E-16). Line of unity (*dark gray*). (B) Percent of genes with outlier expression that are not cancer genes (--, *left*) or are cancer genes (+, right) as per Cancer Gene Census Database, and are not (*gray*) or are (*red*) within +/-150 kb of an HPV integrant (Fisher’s exact, FDR correction, p=2.2E-11 (*left*) and p=7.2E-55 (*right*), respectively. (C) Of 220 genes expressed at outlier levels (Z-score ≥ 2 or ≤-2) in ≥1 tumor and within +/-500 kb of a breakpoint, 16 are cancer genes as shown. *Box and whiskers plot*, Z-scores (*y-axis*) for cancer genes (*x-axis*) in samples harboring nearby breakpoints (*red*) vs. lacking them (*no fill*). (D) Comparison of gene counts (*y-axis*) expressed at various levels (Z-scores, *x-axis*), grouped in 50 kb genomic distances from the nearest breakpoint, in tumors with (*red*) and without (*blue*) breakpoints in those segments. Of 194 genes harboring breakpoints, 155 are expressed as per available RNA-seq data. *Left to right*, breakpoints across the tumors inside or outside genes as indicated; n, counts; q=adj. p-values, binomial test. (E) Percentages of genes expressed at outlier levels (Z ≥ 2) at indicated ploidies (*x-axis*) in absence (*gray*) or presence (*red*) of breakpoints within +/- 500 kb. Ploidy loss, N < 1.5; normal, 1.5 ≤ N ≤ 2.5; gain 2.5 ≤ N ≤ 5; hyper-gain N > 5. *Asterisks*, adj. p<1E-4, binomial test, one-tailed, adjusted by FDR. (F) Q-Q plot of chi-square adjusted p-values calculated from comparison of gene expression Z-score distributions (i.e. sum of the square of Z-scores) at chromosomal loci with HPV-mediated rearrangements (*y-axis*) vs. matched loci without rearrangements (*x-axis*). (G) (*Top*) Depth of sequencing coverage (*y-axis); (bottom*) Z-scores of log2 TPM (*y-axis*) for genes in a tumor with an HPV-linked rearrangement on Chr. 11q13.3 (*x-axis*); genes with outlier expression (*red fill*). Breakpoints (*red vertical lines*) mapping within a 2.4 Mbp region with 8-fold amplification result in outlier expression (Z-score ≥2) for 22 (67%) of 33 genes including *cyclin D1*. Tumor with HPV-linked rearrangement: Z-score <2 (*pink*), outlier Z-score ≥2 (*red*); all other tumors without detectabel(*black*). See also **Table S5**.

Genes with statistical outlier expression levels (Z-scores ≥ 2) are disproportionately higher in frequency within +/- 500 kb of HPV breakpoints compared with those lacking breakpoints, and this difference is greater among cancer genes (Sondka et al. 2018) than non-cancer genes (**Fig. 5B**). Among the 2,898 genes neighboring individual breakpoints across the samples, 220 have outlier expression and 108 are cancer genes (**Table S5.1-2**). Of these, 16 are cancer genes that neighbor an HPV breakpoint and display outlier expression (**Fig. 5C**). Thus, known cancer genes displaying outlier expression in individual tumors are enriched in proximity to HPV integrants, implicating a selective growth advantage imparted by this proximity.

Further analysis of the relationship between a gene’s proximity to an insertional breakpoint and its outlier expression status reveals that integrants’ impacts on gene expression are greatest at distances less than +/- 150 kb (**Fig. 5D**). Outlier levels of gene expression (Z-score ≥ 2) are 6-fold more likely (binomial test, adj. p-value 1.43E-63), and extreme outlier levels of expression (Z-score ≥ 4) are 86-fold more likely (binomial test, adj. p-value 1.23E-86), for genes within +/-150 kb of an HPV breakpoint in affected tumors, compared to the same genes in tumors without those integrants.

Even after accounting for the strong association between HPV integration and genomic ploidy as noted above, outlier expression is significantly enriched in regions with HPV integrants **(Fig. 5E)**. This demonstrates that proximal host gene expression is affected by HPV integrants, even after accounting for effects of differences in local ploidy.

To analyze HPV insertions’ impacts on regional gene expression levels, we defined HPV-linked rearrangements as the broader chromosomal regions containing breakpoints (or clusters), flanked by +/-1 Mbp margins (**Table S5.3**). We compiled 238 HPV-linked rearrangements across the 105 tumors, and compared the cumulative distributions of involved genes’ expression levels in tumors with vs. without such rearrangements in a Q-Q plot, by calculating the sum of the square of their Z-scores. The results show that expression levels of the genes in ~30% of the rearrangements are significantly different from those in the same regions in control tumors without breakpoints (**Fig. 5F**). A representative HPV-linked rearrangement showing expression levels of involved genes reveals that HPV integrants flank and bridge host CNVs and SVs, and induce significant outlier expression of numerous regional genes, in this case including *CCND1* (**Fig. 5G**).

### Diverse forms of genetic disruption by HPV integrants

We investigated relationships between HPV integrants and host genomic features such as annotated genes and regulatory elements. Consistent with previous findings (Bodelon et al. 2016), breakpoints are enriched within fragile sites (binomial test, p=0.0005) and DNase hypersensitive sites (binomial test, p=0.0001), indicating that open chromatin may facilitate HPV integration. Breakpoints are enriched in protein-coding gene promoters (binomial test, adj. p-value 5.49E-8) and exons (binomial test, adj. p-value 4.44E-6), and a slight majority (56.22%) is localized in intragenic regions of annotated genes (**Table S6.1, S6.2**). Breakpoints are enriched in genomic sites bearing marks of active enhancer elements as detected in NHEK cells (e.g. histone 3 lysine 27 acetylation [H3K27ac], binomial test, adj. p=2.35E-6), and excluded from sites with repressive epigenetic marks (e.g. histone 3 lysine 27 methylation [H3k27me3], binomial test, adj. p=8.84E-10). Our findings corroborate associations between HPV integrants and transcriptionally active chromatin bearing H3K27ac marks in HPV-positive cell lines and patient-derived xenografts (Kelley et al. 2017). Moreover, HPV insertions amplified and hijacked a cellular enhancer in subclones derived from W12 cells, a cervical cell line, forming a super-enhancer-like element (Warburton et al. 2018). These data collectively indicate that alterations in host chromatin structure and concomitant changes in gene regulation can serve as a target for and/or result from HPV integration.

Previous analysis of 35 HPV-positive OPSCC identified intragenic HPV integrants which disrupted *RAD51* and *ETS2* (Parfenov et al. 2014). Here, breakpoints map within annotated genes in 79% of tumors with HPV integration (N=194, mean 3.2, median 2, range 1-20 genes per tumor, **Fig. S6.1, Table S6.3**). Intragenic HPV integration coincides with several forms of gene disruption in 71% of these tumors, including CNVs (53%), SVs (41%), and/or virus-host chimeric transcripts (35%). Expression of an additional 79 host genes without detectable intragenic breakpoints is altered by chimeric transcript expression, indicating creation of fusions from nearby HPV insertions. Overall, ~ 92% of tumors harboring HPV integrants have one or more genes (mean 3.97, median 3, range 1-23) disrupted by intragenic insertions and/or chimeric transcript expression.

In 85% of the tumors with HPV integration, a broad range of distinct virus-host chimeric transcripts is detected in RNA-seq data (mean 10.5, median 8, range 1-56 unique transcripts per tumor, **Table S6.4**). We aligned chimeric transcripts expressed in HPV16-positive tumors to the HPV16 reference genome (**Fig. 6A**). A majority of transcripts initiated from viral promoters utilize established viral splice donor sites (ranked by frequency at HPV16 nt. 880 > 226 > 1,302 > 3,632; **Fig. 6A, Fig. S6.2**) spliced to diverse host splice acceptor sites. When aligned to the reference human genome (**Fig. 6B**), chimeric transcripts frequently map in close proximity. These transcripts are related, but vary due to distinct RNA splice site usage. Virus-host splicing junctions frequently map at long distances from insertional breakpoints identified from WGS (**Fig. 6C**). We detect no chimeric transcripts from ~50% of inserted virus sequences as represented by DNA breakpoints. Approximately half of the host genes from which chimeric transcripts are expressed lack intragenic breakpoints. These results show that chimeric transcripts detected by RNA-seq or other RNA-based mapping methods are poor surrogates for detection of all virus-host DNA breakpoints. Since only 56% of virus-host breakpoints are intragenic, whole exome sequencing (WES) also would be a poor proxy for detection of all breakpoints.

**Fig. 6.**
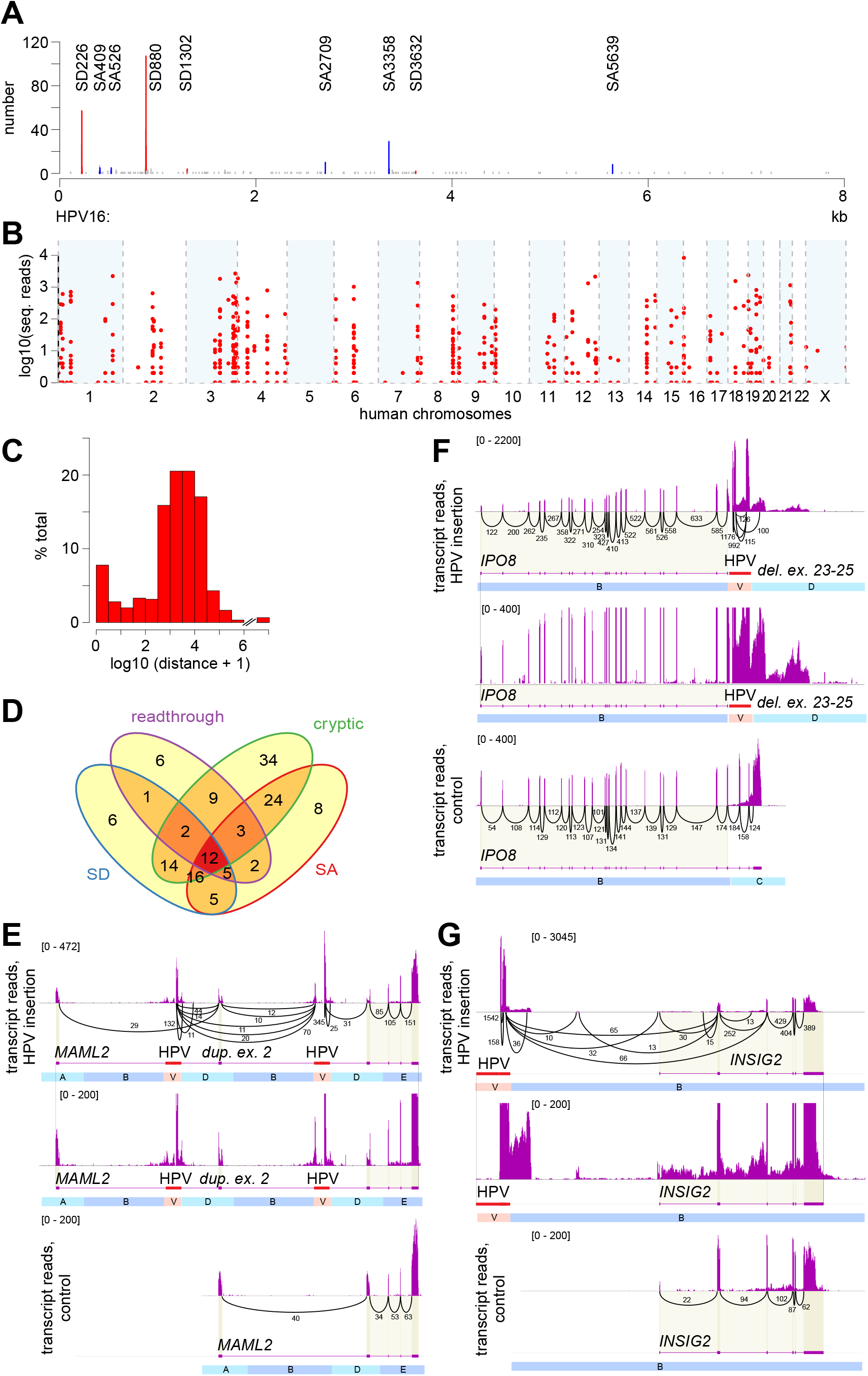
HPV integrants induce various forms of genetic disruption including gene breakage and chimeric transcription. (A) Counts of virus-host chimeric transcript junctions (*y-axis*) in 91 HPV16-positive tumors, aligned to the HPV16 genome (*x-axis*) with known splice donor (SD coordinate, *red*), splice acceptor (SA coordinate, *blue*) and other (*gray*) sites. (B) Counts of split or discordant RNA-seq reads (*y-axis*) in 103 HPV-positive tumors supporting chimeric transcript junctions (n=673), aligned to the human genome (*x-axis*). (C) Frequency distribution (*y-axis*, percent total) of log-transformed genomic distances (*x-axis*) between virus-host junctions from RNA-seq vs. nearest DNA breakpoint (n=604). (D) Venn diagram counts chimeric transcripts expressed at 147 genes, via host splice donor (SD, *blue*, n=61); splice acceptor (SA, *red*, n=75); readthrough transcription (*purple*, n=40) and/or cryptic splice sites (*green*, n=114). (E-G) Sashimi plots depict counts of mapped RNA-seq reads at genes with HPV integrants in affected tumor (*top, center panels*) vs. without integrants in control tumor (*bottom*). HPV sequences not shown to scale. *Center, bottom panels*, identical scale of reads (*y-axis, brackets). Black arcs*, *numbers*, read counts connecting spliced exons. (E) Intragenic HPV integrants in *MAML2* (*red*) flank a ~75 kb duplication including exon 2, and delete small intronic segment C. Gene breaking involves premature transcriptional termination of *MAML2* after exon 2 and de novo initiation of downstream transcripts from HPV. Segment B is truncated for visualization. (F) Intragenic HPV integrants in *IPO8 (red*) delete distal exons 23-25, disrupting 3’ transcripts and upregulating upstream exons. (G) Intergenic HPV integrants upstream of *INSIG2* directly flank a ~700 kb duplication (not shown) on Chr. 2q14. Numerous chimeric transcripts originating from an upstream, intergenic HPV16 integrant are spliced to a novel exon, novel splice acceptor site and exons 1, 2 and 3, causing gene disruption. See also **Table S6** and **Fig. S6**.

Chimeric transcripts are expressed at 147 genes across the tumors, inducing outlier expression of 35% of them (**Table S6.5**). In contrast to canonical splicing of transcripts expressed from amplified host genes (**Fig. 5G**), chimeric transcripts display markedly altered structures when expressed at outlier levels. Chimeric transcripts disrupt the 147 host genes in part via readthrough expression and/or usage of host splice donor, splice acceptor and/or cryptic splice sites (**Fig. 6D**). For example, 31 distinct chimeric transcripts were detected at *CASC8* in a single tumor (**Table S6.5**).

Several forms of genetic disruption are manifested in chimeric transcripts, including expression of novel exons, premature transcriptional termination and gene breaking. Exon-by-exon expression analysis of all host genes affected by fusion transcripts helps visualize these diverse forms of genetic disruption **(Fig. S6.3**). An example of gene breaking (Wheelan et al. 2005), induced by intragenic HPV integrants, was identified at Mastermind-like 2 (*MAML2*), a transcriptional coactivator of Notch (Wu et al. 2002) (**Fig. 6E)**. Premature transcriptional termination of *MAML2* after exon 2 and de novo initiation of downstream transcripts from HPV were observed. Although *NOTCH1* itself is mutated in ~17% of head and neck cancers, Notch pathway signaling is disrupted by other mechanisms including rare driver gene mutations in an estimated 67% (Loganathan et al. 2020). Gene breaking by HPV integrants at *MAML2* uncovers a novel additional mechanism for Notch pathway disruption. Comparable instances of genes harboring intragenic insertional breakpoints are 130-fold more likely to show extreme outlier expression (e.g. Z-scores ≥ 4) compared to the same genes in control tumors lacking such breakpoints (8.4 vs. 0.06%, binomial test, adj. p=1.3E-22). Overexpression of disruptive, chimeric transcripts drives this effect in most cases. In another example, an intragenic HPV integrant within a nuclear importin gene, *IPO8*, results in deletion of 3’ exons 23 to 25 (**Fig. 6F**). Genetic disruption by an intergenic HPV16 integrant upstream of insulin-induced gene 2, *INSIG2*, encoding a negative regulator of cholesterol biosynthesis, is shown in **Fig. 6G**. Here, numerous chimeric transcripts initiated from HPV promoters are spliced to a novel exon, a novel splice acceptor site and exons 1, 2 and 3. Disruption of this HIF1-alpha-inducible gene may facilitate cell growth in a hypoxic tumor microenvironment (Hwang et al. 2017).

### A simple, low-risk HPV69 integrant induces high expression of an imprinted oncogene, *RTL1*

On an infrequent basis, involving ~7% of genes with chimeric transcripts, intragenic HPV integrants markedly upregulate otherwise completely unexpressed genes. Most of these cases affect noncoding RNAs such as *LINC0001* (Zapatka et al. 2020), supporting their putative interactions with HPV (Sharma and Munger 2020) (**Table S7.1**). In a tumor largely devoid of somatic mutations, CNVs and SVs **(Fig. 7A-C)**, a simple insertion of non-oncogenic HPV69 (nt. 2802-2266) was detected in the *DLK1-D103* imprinted domain on Chr. 14q32, without associated CNV or SV. HPV E6, E6*I and E7 transcripts are spliced to exon 1 of retrotransposon-gag-like 1 (*RTL1*), resulting in extreme outlier levels of expression (396 TPM vs. median = 0.011 TPM in controls, **Fig. 7D** and **F**). In the same tumor, another insertion on Chr. 15q21 results in dramatic upregulation of E6/E7 chimeric transcripts and outlier expression of *C15orf65* (27.1 TPM vs. median = 3.44, **Fig. 7E** and **G**). These results demonstrate that even simple integration events by an atypical, likely non-oncogenic HPV type can induce outlier expression levels of candidate driver genes in a primary tumor, very likely contributing strongly to its etiology.

**Fig. 7.**
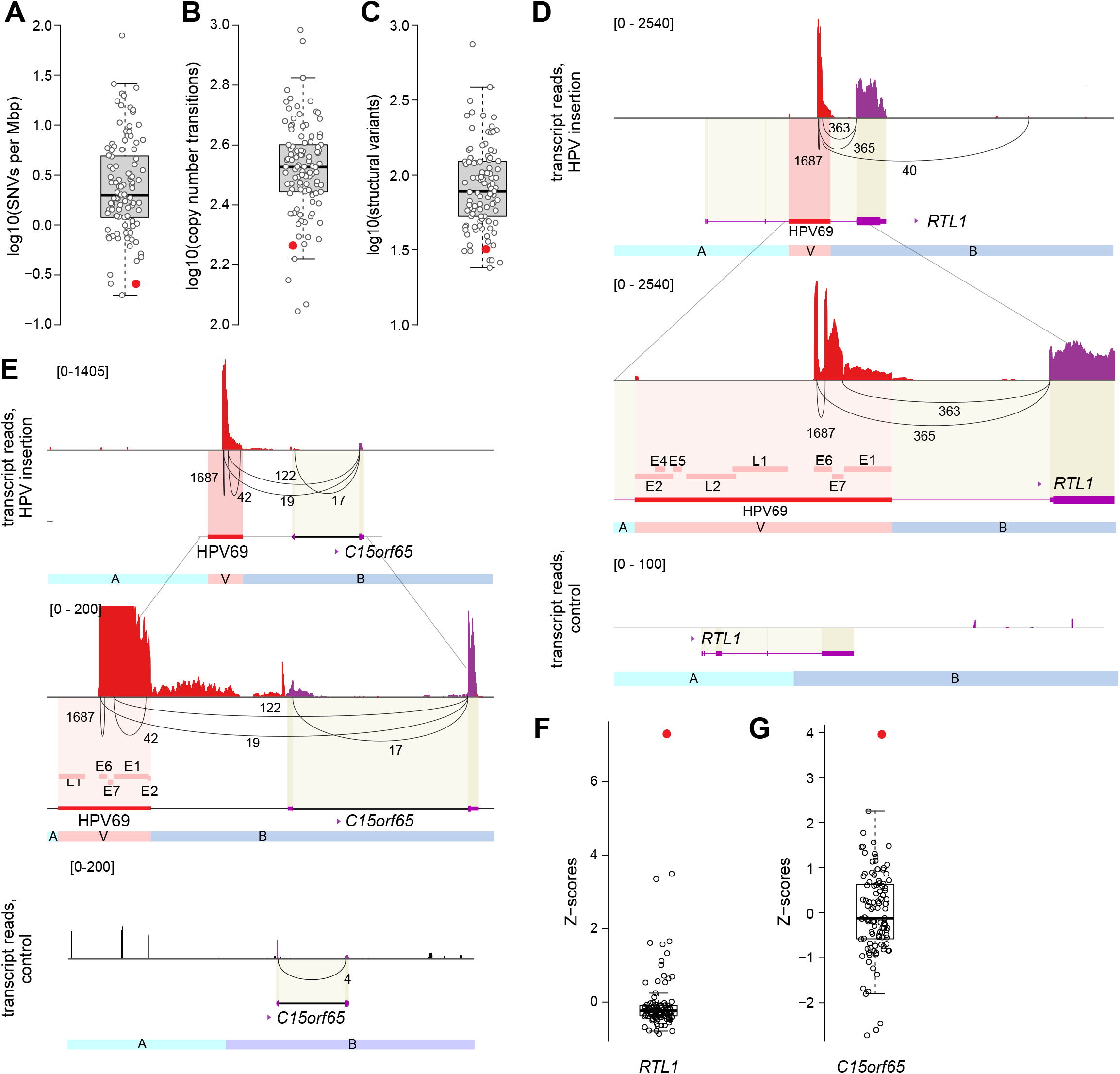
HPV69 integrant induces high expression of an imprinted oncogene, *RTL1, and C15orf65*. (A-C) Box and whisker plots depict log10-transformed counts of (A) SNVs or small indels per megabase pair, (B) copy number step-changes (+/- 0.5N), and (C) SV breakpoints in 105 OPSCC (*circles): red*, tumor with HPV69 integrants; *no fill*, all others. (D, E) Sashimi plots of (*top*) chimeric transcripts initiated in HPV69, spliced to exons of (D) *RTL1* and (E) *C15orf65*, leading to extremely high expression relative to controls. *Black line, numbers*, read counts connecting spliced exons; *bottom*, read counts of conventional transcripts in control tumor. Some *RTL1* fusion transcripts extend past the 3’ transcription termination signal (not shown). (F-G) Box and whiskers plots of expression levels of (F) *RTL1* and (G) *C15orf65* in 103 HPV-positive OPSCC. *Red*, HPV69-positive tumor; *gray*, all others. See also **Table S7**.

In sum, the virus disrupts host transcription in the 95% of the tumors with integration, via intragenic integrants, chimeric transcription, outlier expression, gene breaking and/or de novo expression of noncoding and imprinted genes (**Table S7.2**).

## DISCUSSION

In this analysis, we show that HPV integrants disrupt host genomic structure and expression in essentially all tumors with viral integration. This prospective study was powered to detect significantly recurrent hotspots for integration, and we identified six hotspots near genes that regulate specific biological processes of epithelial stem cell maintenance (i.e. *MYC, FGFR3, SOX2, TP63, KLF5*) and immune cell function (i.e. *CD274*). The pathophysiological significant of this finding is underscored by known roles for these genes in cancer development. Among these are transcription factors known to mediate self-renewal, proliferation and epithelial differentiation and the PDL1 immune checkpoint ligand that facilitates tumor escape from immune surveillance. Second, the numbers of virus-host breakpoints increase directly with the frequency of regional host recombination, supporting a direct role for HPV integrants in SV formation. Third, even after accounting for genomic copy number changes, we find significant enrichment of integrants within 150 kb of genes expressed at outlier levels, thereby implicating HPV integration as the direct cause of genetic disruption. Fourth, essentially all (~95%) tumors with HPV integration have one or more genes disrupted by intragenic insertions and/or chimeric RNA transcription. For example, in a genetically cold tumor lacking frequent SNVs, CNVs and/or SVs, we discover simple HPV integrants inducing outlier levels of chimeric transcripts involving candidate oncogenes. Taken together, our findings indicate HPV integrant mediated host alterations are directly oncogenic.

Three of the recurrent hotspot genes that we find across our large collection of OPSCC samples, i.e. *MYC, TP63 and KLF5*, also have been identified across cervical cancers in a meta-analysis (Bodelon et al. 2016), supporting a common HPV-mediated tumor biology across distinct anatomic sites. The three additional hotspot sites in OPSCC identified here, involving *SOX2, FGFR3* and *CD274*, are previously unreported as hotspots in HPV-positive cancers, although additional cases of HPV integration reported near *CD274* (Koneva et al. 2018) support our findings. Differences by anatomic site could be attributable to limitations in prior sample sizes or methodologies, random fluctuations between sample collections, or specific biological differences across the tissue sites of HPV infection. The latter explanation may underlie why we did not find the *RAD51B* or other hotspots in OPSCC as reported in cervical cancers (Ojesina et al. 2014; Bodelon et al. 2016; Cancer Genome Atlas Research et al. 2017). We note that studies utilizing WES miss half of HPV integration breakpoints. RNA-seq can detect chimeric transcripts but it mismaps the corresponding integrant templates by up to several megabasepairs. Moreover, RNA-seq altogether misses the half of genomic integrants that we find do not generate chimeric transcripts. In the 874 integration breakpoints identified here, we find no instances of identical breakpoints shared across any two independent tumors (Hu et al. 2015; Dyer et al. 2016), highlighting the robustness of our approach.

Here we show that HPV integration disrupts host genes by mechanisms including genomic amplifications, deletions, inversions, translocations and other rearrangements, intragenic insertion, initiation of chimeric transcripts from heterologous viral promoters, and introduction of promoter, splice acceptor, splice donor and transcriptional terminator elements. Each of these disruptive mechanisms can lead to alterations in gene expression, structure and function. HPV integrants amplified oncogenes including *MYC, CCND1*, and *SOX2*, and an immune checkpoint ligand *CD274;* disrupted known tumor suppressors including *MAML2, EP300* and *INSIG2;* and induced aberrant overexpression of the imprinted oncogene *RTL1*. Our data indicate that HPV-mediated insertional mutagenesis is at least as disruptive of host genomic homeostasis as are SNVs, but would not be detected or resolved by widely utilized platforms such as WES or targeted cancer gene panels (Chung et al. 2015). We conclude that this extensive disruption of host genomic structures and functions we report here implicates HPV insertional mutagenesis as a promoter of the malignant phenotype.

We note that the primary HPV-positive OPSCCs studied here could not be further manipulated experimentally. However, prior studies of unrelated HPV-positive cancer cell lines demonstrated that their proliferation and viability depend on genetic disruptions caused by HPV integrants (Akagi et al. 2014; Shen et al. 2017; Warburton et al. 2018; Broutian et al. 2020). The best-studied of these is the HeLa cell line, established in culture from Henrietta Lacks’ aggressive cervical cancer. The key instigator of tumorigenesis in this case consists of the HPV-mediated amplifications, rearrangements and long-range interactions with enhancer elements that induce massive upregulation of *MYC* (Adey et al. 2013; Shen et al. 2017). Experimental deletion of these HPV integrants resulted in marked reductions in *MYC* expression (Shen et al. 2017). Similarly, we demonstrated collaboratively that *MYC* overexpression drives proliferation of the cervical neuroendocrine cell line GUMC-395, in which HPV integrants directly flank a ~40-fold amplification of *MYC* (Yuan et al. 2017). *MYC* overexpression drives tumorigenesis upon *TP53* loss (Zindy et al. 1998). Documented cooperative interactions between HPV E6 and MYC in inducing telomerase (*hTERT*) expression and keratinocyte immortalization further support the pathophysiological significance of viral integration at this hotspot (Zhang et al. 2017). Moreover, we have shown that genetic knockdown or small molecule inhibition of PIM1 induces cell death in UPCI:SCC090, a head and neck cancer cell line in which HPV integrants directly flank a 16-fold amplification of *PIM1* (Broutian et al. 2020). These archetypal, experimentally tractable examples each illustrate the contributions of HPV integrant-mediated transcriptional alterations to cellular transformation and the malignant phenotype.

A “genetically cold” tumor with very low frequencies of somatic SNVs, CNVs and SVs (**Fig. 7**) harbors two simple integrants of HPV type 69 on Chrs. 14 and 15. Highly expressed chimeric transcripts are initiated from both integrated HPV promoters and include the viral E6/E7 genes. The Chr. 14 transcripts are spliced to exons of *RTL1*, an imprinted oncogene, while Chr. 15 transcripts are spliced to *C15orf65* exons, encoding a protein of unknown function. Mouse models of hepatocellular carcinogenesis involving an engineered Sleeping Beauty transposon showed highly analogous, frequent activating insertions upstream of *RTL1*, resulting in overexpression of transposon-*RTL1* chimeric transcripts (Riordan et al. 2013). Overexpression of *RTL1* in adult mouse liver led to tumor formation in 86%, revealing that it is a potent oncogene. Aberrant expression of *RTL1* also has been reported in melanoma (Fan et al. 2017). By analogy, we conclude that marked upregulation of these two genes due to simple HPV integrants (without associated CNVs or SVs) also could drive OPSCC tumorigenesis directly, drawing parallels to retrovirus and activating transposon integrants causing cancer in numerous contexts (Kawakami et al. 2017; Bushman 2020).

Persistent viral infection and E6 and E7 expression initiate carcinogenesis and promote genomic instability, but additional genetic events are necessary for cancer development. Persistent infection comprises stable maintenance of HPV episomes, initiated years to decades prior to cancer diagnosis, but the timing of viral integration in tumorigenesis remains uncertain. HPV integration increases in frequency with severity of cervical dysplasia, and is present in the majority of cervical cancers (Bodelon et al. 2016), suggesting that it drives clonal selection and carcinogenesis. In vitro models demonstrated that HPV integration confers a selective growth advantage, attributed to increased expression and stabilization of viral transcripts encoding oncoproteins (Jeon et al. 1995).

Our working model holds that initial HPV integration occurs randomly but preferentially in genomic regions of open chromatin. Simple HPV integration may occur at sites of individual DNA double strand breaks. By contrast, at sites of multiple such breaks, HPV may capture transient free ends of intervening host DNA, generate virus-host concatemers via rolling circle amplification typically initiated from the viral origin of replication, and after recombination and repair, result in clusters of integrants, CNVs and SVs (Akagi et al. 2014). Resulting, viral-host concatenated arrays at insertion sites occur because viral replication depends upon host DNA damage response pathways (Gillespie et al. 2012; Reinson et al. 2013). Individual cell clones with growth advantages imparted by HPV integration are positively selected, resulting in associations between insertional breakpoints and CNVs, SVs, cancer genes and outlier levels of expression specific to each emerging cancer. We formulated this mechanistic model, termed the HPV-mediated looping model, to explain the formation and impacts of HPV integrants in cultured cancer cells (Akagi et al. 2014). Here we have provided comprehensive additional evidence from primary cancers to support and extend this model.

Limitations to our approach include a lack of analysis of HPV insertion-mediated effects on small RNAs, long noncoding RNAs, epigenetic controls or chromatin changes. We also have not fully resolved the genomic architecture of some of the most complex HPV-mediated SVs, so currently we are utilizing complementary methods including long-read DNA sequencing to address this. We plan to investigate tumor heterogeneity of HPV integration-mediated SVs, CNVs and chimeric transcription at subclonal or single cell levels.

We conclude that host genomic alterations and gene disruption induced by virus integration are necessary driver events in a high proportion of HPV-positive cancers. We acknowledge that approximately 23% of tumors have episomal HPV DNA only, implicating alternative mechanisms of cancer promotion not mediated by viral integration. Although absence of HPV breakpoints indicate exclusively episomal virus in some tumors, WGS short reads cannot discriminate exclusively integrated from concurrent presence of episomes. HPV insertion-mediated CNVs and SVs in primary tumors are consistent with our looping model based on cell line data (Akagi et al. 2014), in which transient virus-host circular intermediate structures are amplified, recombined and repaired with flanking host sequences. We and others have reported similar impacts in tumors caused by Merkel cell polyomavirus (Starrett et al. 2020) and hepatitis B virus (Jiang et al. 2012). Thus, growing evidence supports impacts of virus-mediated insertional mutagenesis with genomic instability and genetic disruption as a common hallmark of the ~10% of cancers caused by DNA viruses.

## METHODS

### Study population

Patients with newly diagnosed, HPV-positive OPSCC, presenting at Ohio State University Comprehensive Cancer Center from 2011-2016, provided written, informed consent for genomics studies and prospective collection of clinical data (Gillison et al. 2019). This study was approved by Institutional Review Boards at Ohio State University (OSU) and the University of Texas MD Anderson Cancer Center (MDACC). The overall study population included 105 patients with HPV-positive OPSCC, including 86 from OSU and 19 studied by The Cancer Genome Atlas project (TCGA, https://gdc.cancer.gov/). Clinical characteristics were reported previously (Gillison et al. 2019). A statistical power calculation indicated that at least 96 tumors would be required to provide 90% power to detect recurrent integration events within the same 1 Mbp window in 4% of tumors. An additional 53 deidentified, fresh-frozen OPSCCs were obtained from an MDACC head and neck cancer specimen bank, to increase genomic DNA sample numbers to a total of 158 HPV-positive OPSCC for detection of additional recurrent integration hotspots. All tumors were confirmed p16-positive by immunohistochemistry and HPV-positive by quantitative PCR (Gillison et al. 2019).

### Genomic DNA WGS analysis

OPSCC tumors were snap-frozen and microdissected to ensure >70% tumor content. Genomic DNA was isolated from paired tumors and matched normal blood leukocytes. Tumors’ HPV status and genomic DNA sample quality were measured (Gillison et al. 2019). In the OSU tumors, 34 HPV-positive OPSCC T/N pairs were sequenced at ~90x mean depth of coverage by Complete Genomics (CGI) WGS (Gillison et al. 2019). The CGI aligner was used to map paired-end WGS reads against the human reference genome assembly GRCh37 (hg19) (Carnevali et al. 2012). Illumina WGS data from 52 OSU HPV-positive OPSCC T/N pairs were generated at New York Genome Center (NYGC), including 40x mean depth of coverage for normal samples and ~90x for tumors. Illumina WGS data for 19 HPV-positive cancers were downloaded from TCGA. GATK v.3 was used to identify duplicate reads, realign reads surrounding indels, and recalculate alignment quality scores (McKenna et al. 2010). To estimate HPV copy numbers per sample, mean depth of sequencing coverage in the viral genome was divided by mean depth of autosomal coverage. HPV copy number was compared in tumors with and without HPV integrants by t-test.

### HPV insertional breakpoint detection

To detect virus-host breakpoints, raw Illumina sequence reads were aligned by BWA MEM (v0.7.15) (Li and Durbin 2009) against a hybrid reference assembly combining human (hg19) and 15 high-risk HPV genomes (hereafter named hg19+HPV). The virus types and GenBank accession numbers were HPV16 [NC_001526.2]; HPV18 [NC_001357.1]; HPV31 [HQ537687.1]; HPV33 [HQ537707.1]; HPV35 [M74117.1], HPV39 [M62849.1], HPV45 [X74479.1], HPV51 [M62877.1], HPV52 [X74481.1], HPV56 [X74483.1], HPV58 [D90400.1], HPV59 [X77858.1], HPV66 [U31794.1], HPV68 [DQ080079.1] and HPV69 [AB027020.1]). Discordant read pairs and split reads supporting the presence of virus-host breakpoints were extracted using three breakpoint callers, i.e. Hydra (Quinlan et al. 2010), Delly (Rausch et al. 2012), and SplazerS (Emde et al. 2012). HPV breakpoints supported by at least two independent discordant or split read pairs by each of at least two callers or by four or more pairs by at least one caller were analyzed further. Discordant pairs were extracted from CGI data using custom scripts with individual hybrid genome reference assemblies. Virushost breakpoints were called based on five or more supporting pairs.

### Confirmation of HPV insertional breakpoints by HPV capture-seq

Custom Agilent SureSelect baits were designed to capture HPV (Warburton et al. 2018). Genomic DNA of selected tumors (i.e. GS18109, GS18041, GS18006) was hybridized and libraries were prepared following the manufacturer’s directions. We generated 20 to 50 million read pairs per sample. HPV breakpoints were detected using published methods (Quinlan et al. 2010). In a separate approach, ~10% of breakpoints identified from WGS data were randomly selected for confirmation by Sanger sequencing. Custom PCR primers were designed to amplify across each breakpoint (Akagi et al. 2014). PCR products were sequenced using conventional Sanger sequencing. If unsuccessful, PCR products were cloned using the TOPO TA cloning kit (Life Technologies) and sequenced bidirectionally using M13F forward and M13 reverse primers.

### Statistical analysis of demographic characteristics and survival outcomes

Characteristics of patients with exclusively episomal vs. integrated HPV were compared by t-test or chi-squared test as appropriate. Progression-free survival, defined as date of diagnosis to cancer progression or death, was compared in tumors with exclusively episomal vs. integrated HPV by Kaplan-Meier analysis and were compared with a log-rank test.

### Hotspot detection

Identification of recurrent hotspots of HPV integration across tumors could be confounded by breakpoints clustered in individual tumors, which are not mutually independent. Therefore, breakpoint clusters within 500kb genomic segments were identified in each tumor using mergeBed function of Bedtools (Quinlan and Hall 2010), and particular breakpoints supported by the highest read counts were identified as representative breakpoints. We identified 238 representative breakpoints in 105 OPSCC studied by WGS, and 80 more in 53 OPSCC studied by HPV capture-seq (**Table S3-4**). To identify expected breakpoints, we performed in silico simulations to calculate the probability of observing loci with recurrent integration. The human genome was segmented into ~3,000 independent, non-overlapping, 1 Mbp tiles. We simulated 238 or 318 representative breakpoints hitting these tiles 1 million times. The expected distribution of HPV breakpoints would be random, allowing us to calculate empirical probabilities of the observed distribution.

Because annotated genes themselves are not randomly distributed in the human genome, we used a second, gene-centric approach to identify genes that are recurrently targeted by HPV integration. To calculate the expected number of times genes would be hit within 500 kb by simulated integrants, two million in silico random integration sites were generated. The observed number of hits per gene for 238 representative HPV breakpoints as identified in WGS data was compared to this expected distribution. Genes with more breakpoints than expected by chance were selected as hotspots for integration (one-tailed binomial test, adj. p<0.01).

### Gene ontology

To account for bias introduced by clustered virus-host breakpoints, simulations were run with 238 representative breakpoints. As a control, 2 million random breakpoints were generated *in silico*. Gene models from Gencode v18 were used to identify ≤4 nearest neighbor genes located within 500 kb from each HPV breakpoint and each control breakpoint. Genes were annotated as per Gene Ontology Resource (www.geneontology.org). Binomial test was used to compare the observed versus expected frequencies of breakpoints found within 500 kb of genes in the Biological Process category (one-tailed test). P-values for GO terms with ≥3 virus-host breakpoints were calculated. P-values were adjusted by FDR multiple testing correction.

### Detection of small variants, SVs and CNVs

Illumina WGS reads were aligned against human reference genome hg19 using BWA.aln version 0.7.15 (Li and Durbin 2009). To detect SNVs and small indels, we used variant caller packages Mutect (Cibulskis et al. 2013), LoFreq (Wilm et al. 2012) and Strelka (Saunders et al. 2012). Variants supported by two or more callers were selected for further analysis (Gillison et al. 2019). For CGI WGS data, small variants were called using the CGI Cancer Genomics pipeline at default settings. CNVs were detected by comparing the depths of coverage in WGS data in T/N pairs using CNANorm with smoothening and segmentation (Gusnanto et al. 2012). Ratios of sequencing depth of coverage were calculated for 2 kb bins in each tumor vs. its matched normal sample and were smoothed using the DNAcopy algorithm within CNANorm. To visualize sequence alignments, depth of coverage, discordant pairs, and breakpoints, we used Broad Institute’s Integrative Genome Viewer (IGV) (Robinson et al. 2011). Structural variants were identified using three callers, i.e. Crest (Wang et al. 2011), Delly (Rausch et al. 2012), and BreakDancer (Chen et al. 2009). SVs identified by two or more callers were selected for further analysis.

### Relationships between HPV breakpoints, CNVs and SVs

To evaluate relationships between HPV breakpoints and CNVs, first we defined 100 kb bins across the human genome. Estimated ploidies for each bin were determined by identifying the majority ploidy segment representing the bin (Gusnanto et al. 2012). Bins with majority ploidy N≥2.5 were defined as copy number amplifications, while bins with majority ploidy N≤1.5 were defined as copy number losses. Hyperamplified bins were defined by majority ploidy N≥ 4 or 5 depending on analysis.

Using CNANorm outputs of segments and ploidies, we identified copy number transition sites as boundaries between 100 kb segments with ploidy changes of ≥0.5N. Genomic copy numbers and copy number transitions were compared between bins with HPV breakpoints vs. those without. Associations were evaluated using a one-tailed binomial test. *P*-values were adjusted by FDR multiple testing correction. To evaluate associations between HPV breakpoints and SVs (i.e. chromosomal translocations, deletions, inversions, rearrangements), we conducted a similar analysis comparing bin lengths of 100, 200 and 500 kb genomewide. Statistical analysis (as described above) was performed using Bioconductor (http://www.bioconductor.org) packages.

### Linked-read sequencing

Linked-read genomic DNA libraries were prepared from selected tumors using the 10x Genomics Chromium platform, and 2×150 bp paired end sequences were generated at approximately 30-40x depth of coverage. Data were aligned to the hg19+HPV hybrid reference assembly, and mapped read barcodes were identified using LongRanger v. 2.2.2 (10X Genomics). To resolve haplotypes at particular loci, germline SNVs were identified from WGS data of matched controls. Loupe (10X Genomics) and custom scripts using R graphics were used to visualize alignments and haplotypes. The fraction of distinct haplotype-specific barcodes mapping in each 2 kb genomic segment at the *CD274* locus of GS18047 that also was shared with HPV was calculated.

### RNA-seq libraries and analysis

Total RNA was isolated and RNA-seq was performed as described (Gillison et al. 2019). RNA-seq reads from HPV-positive OPSCCs were aligned against the human GRCh37 reference using STAR aligner version 2.4.2a (Dobin et al. 2013). RNA-seq reads from all TCGA samples also were similarly aligned. To determine expression levels of each transcript, transcript structures first were downloaded from Gencode v.18 (http://www.gencodegenes.org/) as gene models. Aligned reads were counted using featureCounts (Liao et al. 2014). Raw counts were transformed into transcripts per million reads (TPM) using gene lengths defined by the union of all annotated exons. Genes with low expression values (maximum TPM across all samples < 2) were excluded from further analysis. A pseudo-count of one was added to all TPM values to avoid undefined data upon log transformation. Resulting log2 TPM values were normalized and batch-corrected using Bioconductor sva function ComBat (Leek et al. 2012).

### RNA expression analysis (Z-scores)

Gene-level expression data (log2-transformed TPM values) for 103 samples with available RNA-seq data were analyzed. For each annotated gene, Z-scores were calculated by standardizing the data based on the mean and standard deviation across the samples.

Fractions of genes with outlier expression (absolute value of Z-score ≥ 2) were compared in cases with vs. without HPV breakpoints nearby (i.e. +/- 500 kb). The statistical significance of this enrichment in outlier gene expression in genes with HPV breakpoints was assessed using one-tailed binomial test. P-values were adjusted by FDR multiple testing correction. We also calculated the sum of the square of Z-scores for all genes within HPV-linked rearrangements (HPV breakpoint clusters +/- 1 Mbp margins). Under the null hypothesis of no effect, the sum of the squared Z-scores would be expected to follow a chi-squared distribution with the number of degrees of freedom equal to the number of genes. Using this chi-squared distribution, these summary measurements were converted to p-values in order to compare regions containing different numbers of genes. We plotted the distribution of the sum of squared Z-scores using quintile-quintile (Q-Q) plots.

### Relationships between HPV breakpoints and annotated host genes

Genes containing intragenic HPV breakpoints were identified using GenCode database (v18). Overlaps between breakpoints and chromatin marks as reported by the ENCODE project were identified using the UCSC genome browser (genome.ucsc.edu). The observed distribution of HPV breakpoints was compared to 2 million random breakpoints generated in simulations as a control. Comparisons were performed using a binomial test. *P*-values were adjusted by FDR multiple testing correction.

### Chimeric HPV-host transcript analysis

RNA-seq reads were aligned against hg19+HPV reference assembly using tophat-fusion (Kim and Salzberg 2011). RNA-seq reads were aligned against custom virus-host chimeric models using GSNAP (Wu et al. 2016). At least one split read and ≥2 discordant pairs were required to identify chimeric HPV-host transcripts, which were visualized using the Sashimi plot function of IGV (Katz et al. 2015). To confirm structures of selected chimeric transcripts, de novo RNA-seq assembly was performed using Trinity (Grabherr et al. 2011).

### Detection of HPV-mediated intragenic expression changes

Exon start and stop coordinates were downloaded from GenCode v18. Sums of read basepair counts mapping to each exon were counted for each sample using Samtools bedcov function (Li and Durbin 2009). Basepair coverage was normalized by RNA-seq library size. Batch correction was performed using SVA ComBat (Leek et al. 2012). Mean depths of coverage were calculated on an exon-by-exon basis for each sample, i.e. for exons before and after HPV chimeric transcript junctions. Ratios calculated from after coverage vs. before coverage read counts were log2-transformed for each sample, and Z-scores were calculated across all samples.

## DATA AND REAGENT ACCESS

Original source WGS and RNA-Seq data have been deposited at the European Genome-phenome Archive (EGA; http://www.ebi.ac.uk/ega/), hosted by the European Bioinformatics Institute (EBI), European Molecular Biology Laboratory (EMBL), under accession numbers EGAS00001002393, EGAS00001003228 and EGAS00001003237. We did not generate new code or unique reagents in this study.

## ACKNOWLEDGMENTS

We gratefully acknowledge the participation of patients with oropharyngeal cancers at Ohio State University (OSU) who enrolled in our research study; members of the Gillison and Symer laboratories for insightful comments at various stages of this study; Elisa Venturini, Karen Bunting, Benjamin Hubert and Dayna M. Oschwald for expert help with project management at New York Genome Center; the Genomics Shared Resource at OSU Comprehensive Cancer Center (OSUCCC) for DNA and RNA quality assays; and Jordan Pietz (MDACC) for help preparing graphical figures. This study was funded by the Oral Cancer Foundation (MLG); OSUCCC (MLG, DES); University of Texas MDACC (MLG, DES); Ohio Supercomputer Center (PAS0425, DES); Ohio Cancer Research Associate grant (GRT00024299, KA); Cancer Prevention Research Institute of Texas (CPRIT, MLG); National Cancer Institute grant R50CA211533 (KA). Dr. Gillison is a CPRIT Scholar in Cancer Research.

D.E.S. would like to dedicate this study to the memory of his father, Donald G. Symer, who died of COVID-19 during the preparation of this manuscript.

## AUTHOR CONTRIBUTIONS

Conceptualization, MLG, DES; Methodology, MLG, DES, KA, HMG, KRC, AE, BS, MZ, NR; Formal Analysis, MLG, DES, KA, HMG, KRC, AE, BS, MZ, ZD, NR; Investigation, WX, BJ, YS, GL; Resources, AA, EO, AE; Data Curation, KA, HMG, AE, NR, BS, JS; Writing-Original Draft, MLG; Writing-Review & Editing, MLG, DES; Supervision, MLG, DES; Funding Acquisition, MLG.

